# Reconciling the *clk-1* and aging paradox and categorizing lifespan curves by taking individual specificity into account

**DOI:** 10.1101/213512

**Authors:** Yaguang Ren, Congjie Zhang, Wenxuan Guo, Chao Zhang

## Abstract

The *clk-1* gene encodes the demethoxyubiquinone (DMQ) hydroxylase that is required for biosynthesis of ubiquinone (coenzyme Q). Deletion of *clk-1* was lethal in mice, and its mutation in *C. elegans* mildly extended lifespan, slowed physiological rate and led to sickness. We found that if growth retardation was taken into account the average lifespan of *clk-1* mutants would not be prolonged or would be shortened. In addition, recent study showed that knocking down of *clk-1* shortened lifespan. Although the extension of lifespan in *clk-1* mutants was mild and was not observed sometimes, some progenies indeed had prolonged maximum lifespan even if retardation of growth was taking into account. These paradoxes implicate the existence of individual specificity in the aging process even in the same cohort, just like a drug is beneficial for some people while for others it is detrimental. We further categorized lifespan curves into five kinds of patterns according to the lifespan alternations observed in organisms: N (normal); L (long-lived); S (short-lived); F (flattened); ST (steepened), and found that the curve of *clk-1* mutants fit into the F pattern. The reasons behind the individual specificity and its implications in aging process deserves further investigations.

## 1. Introduction

Coenzyme Q (CoQ), also known as ubiquinone (UQ), is mainly localized in mitochondrion and functions to transport electrons in the respiratory chain (Larsen and Clarke 2002; Vajo et al., 1999). *Clk-1* encodes the *C. elegans* ortholog of COQ7/CAT5 that is necessary for biosynthesis of Coenzyme Q (Ewbank et al., 1997; Gu et al., 2017). Mutation of *clk-1* led to pleiotropic phenotypes including retardation of growth, reduced brood size, long defecation cycle, and mild extension of lifespan (Lakowski and Hekimi 1996; Takahashi et al., 2012). *Clk-1* was one of the first genes found to be correlated with aging and was widely used to explore aging and neurodegenerative disease related mechanisms (Gu et al., 2017; Lakowski and Hekimi 1996; Larsen and Clarke 2002). However, RNAi knocking down of *clk-1* shortened lifespan, although growth rate was not slowed down (Ren et al., 2015; Zhang et al., 2017). In mice, deletion of *clk-1* was lethal and its decreased expression promoted neuroinflammation and subsequent death of dopaminergic cells (Gu et al., 2017; Nakai et al., 2001). These results imply the complication of the effects of *clk-1* deficiency on aging.

## 2. Material and methods

### 2.1 *C. elegans* strains

The wild type strain N2 and the *clk-1* mutant MQ130 *clk-1(qm30) III* were obtained from Caenorhabditis Genetics Center (University of Minnesota, USA), and were grown on E. coli OP50 seeded NGM plates as described by Brenner (Brenner 1974).

### 2.2 Lifespan tests

Lifespan was tested as described with minor modifications (Ren et al., 2015). Briefly, twenty gravid adults were transferred onto OP50 seeded NGM plates, and were removed after laying eggs for eight hours. When the F1 progenies reached L4 stage, transferred one hundred of them onto freshly prepared NGM plates every day during the reproduction period, and every two or three days after egg laying ceased. Worms that did not respond to gentle nose touch with the toothpick were scored as dead, and that crawled off the plates or died from egg hatching in the uterus were censored.

### 2.3 Phenotype analysis

Phenotypes including body size and brood size were analyzed as described (Ren et al., 2012).

### 2.4 Statistical analysis

The Kaplan-Meier method and log-rank test were performed for analysis of lifespan data. Paired student’s-t test was used for growth and reproduction analysis. Differences in means of data were considered statistically significant at p < 0.05. Data were analyzed using the Graphpad Prism 5.0 software.

## 3. Results and discussion

Consistent with previous studies, we found phenotypes including slowing down of growth (Fig 1A, B), decreased fertility (Fig 1C), retardation of reproduction (Fig 1C), and mild extension of lifespan in worms with *clk-1* mutation (Zhang et al., 2017). But the lifespan extension was not reproduced quite well (Fig 1D), perhaps due to its relatively minor effect on aging. Interestingly, the average lifespan of worms with *clk-1* mutations was even found to be shortened (Larsen and Clarke 2002), although other labs reported mild extension by around 5% to 20% or 1 to 3 days (Lakowski and Hekimi 1996; Takahashi et al., 2012). The post growth span (scored from the time when rapid growth ceased) of the *clk-1* mutants was not prolonged (Zhang et al., 2017), and was even shortened as shown here (Fig. 1E), suggesting retardation of growth contributed largely if not completely to lifespan extension resulted from *clk-1* mutation.

**Fig. 1.**
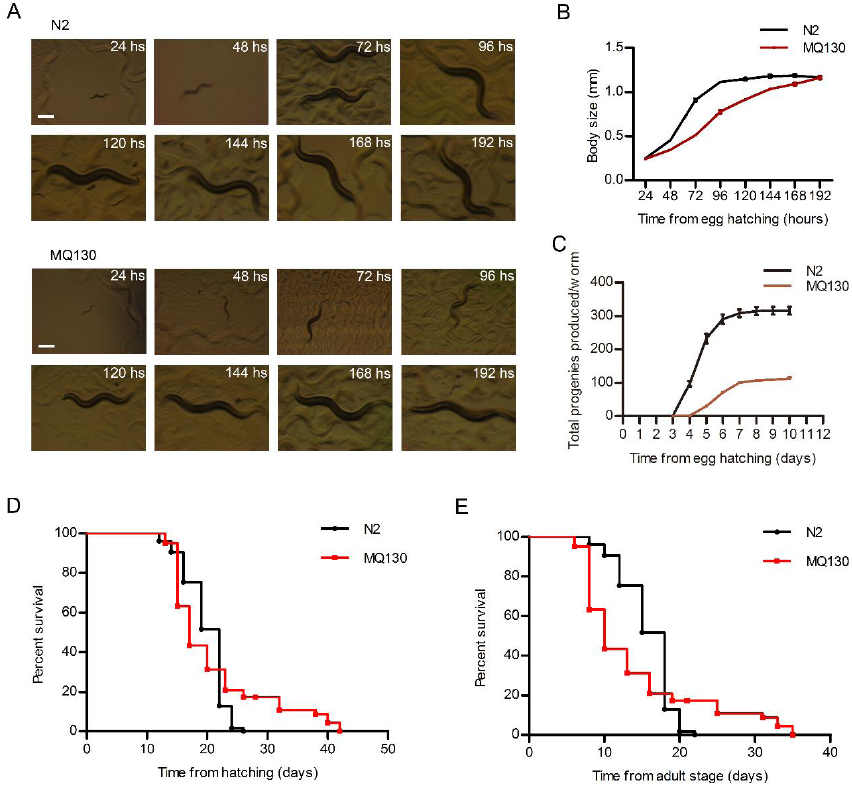
Average lifespan of the *clk-1* mutants (MQ130) was reduced when retardation of physiological behavior was taken into account. (A, B) Growth of *clk-1* mutant strain MQ130 was retarded comparing to that of wild type N2, p<0.05. Scale bars= 200μm. (C) Reproduction was reduced and egg laying was retarded in *clk-1* mutants, p<0.01. (D) Average lifespan of MQ130 (20.28±7.76 days, n=61) was similar to that of N2 (19.60±3.40 days, n=75), p>0.05. (E) Average adult lifespan of MQ130 (13.28±7.76, n=75) was reduced comparing to that of N2 (15.60±3.40, n=61), p<0.05.

The physiological behavior was also retarded in adults (Vajo et al., 1999), which might also contribute to lifespan extension. Then is it logical to interpret slowing down of physiological behavior as slowing down of aging? We speculate that the answer should be “not”, considering retardation of physiological behaviors such as growth is quite common in *C. elegans* under detrimental conditions. For example, in large-scale genomic RNAi screens growth was found retarded by knocking down hundreds of genes respectively (Simmer et al., 2003). Comparing to vertebrates, worms’ growth was much easier to be affected by environmental factors such as perturbation of temperatures (Hirsh and Vanderslice 1976). If slowing down of growth is interpreted as slowing down of aging, the aging related factors will be too many. Thus, sometimes the extension of the post-growth span rather than the entire lifespan should be considered as evidence for slowing down of aging. Although the average lifespan was not prolonged by *clk-1* mutation in some cases, the maximum lifespan was increased and the minimum lifespan was decreased as shown by crossing of survival curves of the MQ130 and N2 strains. This phenomenon was also shown in previous studies (Larsen and Clarke 2002; Zhang et al., 2017). The maximum lifespan of the *clk-1* mutant was more than 40 days, which cannot be explained solely by retardation of growth. The results suggest there is individual specificity in aging process, which may be existed in all organisms. For example, most people die at around seventy years old, while a few can live up to more than one hundred years. It is reasonable that some worms were unusually long-lived while some others were unusually short lived by loss of function of *clk-1*. Just like one drug is beneficial for some people, while for others it is detrimental.

According to the excessive response model, when the prooxidant stress goes high the antioxidant stress will go higher and lower ROS will be observed in the long-term (Ren and Zhang 2017). This strategy should be used to fight against more serious and unpredictable conditions forthcoming (Ren and Zhang 2017). In worms with mitochondrial dysfunctions, retrograde responses were indeed motivated including increased expression of antioxidant enzymes and chaperones, increased autophagy, reprogramming of energy metabolism, and increased activity of the hypoxia-inducible factor HIF-1 (Shore et al., 2012). Some of these responses were reported to have anti-aging effects besides enhancing adaptions (Shore et al., 2012). Lifespan should be affected synthetically by pro-aging factors such as the side effects of cellular dysfunctions, and the anti-aging factors such as some of the retrograde responses (Fig. 2A). Due to individual specificity, in some of the *clk-1* mutated worms the effects of the pro-aging factors may overwhelm that of the anti-aging factors. Thus, lifespan would be shortened. But in other worms the effects of anti-aging factors may overwhelm that of the pro-aging factors, and the lifespan should be prolonged. This idea explains why the survival curve of *clk-1* mutants was flattened and was crossed with that of N2, as was shown here and in previous studies (Larsen and Clarke 2002). Here, the lifespan curves were classified into five kinds of patterns (Fig. 2B).

**Fig. 2.**
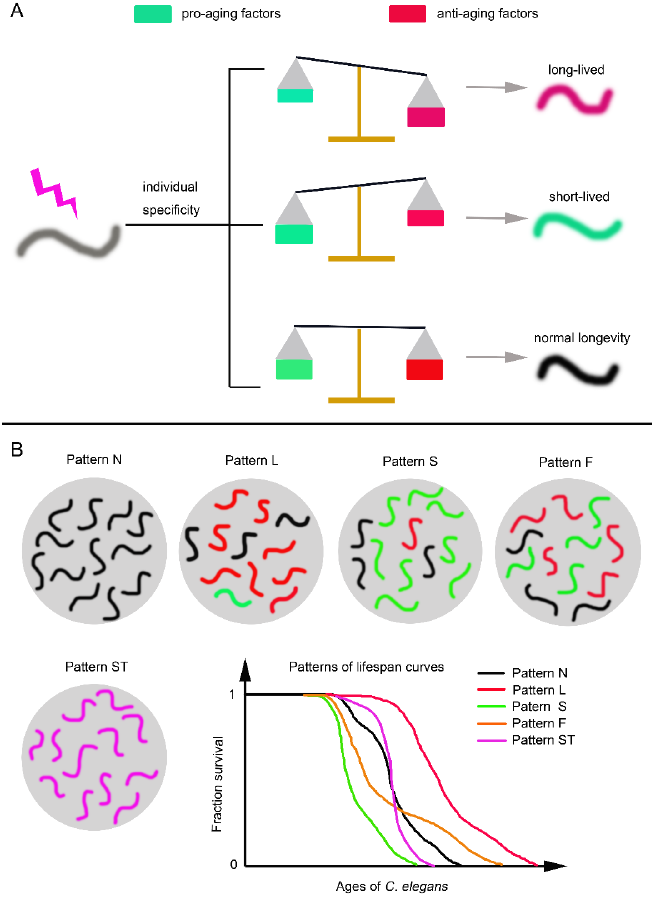
Categorization of lifespan curves by taking individual specificity into account. (A) Under stressful conditions, side effects and retrograde responses will arise. But there are individual specificity in both aspects. As the result, many worms’ individual lifespan will be different from each other. (B) Categorization of lifespan curves based on differences in individual specificity. Black, red, and green colors indicate worms that have relatively normal, prolonged, and shortened lifespan respectively. Purple color indicates cohort of worms whose variations of individual lifespan are smaller than that of N2.

The curve that is similar to that of *clk-1* mutant worms is defined as F (flattened) patterned. Due to residual levels of *clk-1* mRNA, in *clk-1* RNAi worms the less severe side effects may activate lower extent of retrograde responses. It is likely that in *clk-1* RNAi worms the anti-aging factors might fail to compete with the pro-aging factors, and led to decreased average lifespan. The lifespan curve of *clk-1* RNAi worms thus fits into the S (short-lived) pattern. Worms with mutations in *mev-1*, the gene that encodes subunit of mitochondrial respiratory chain complex II, should have S patterned survival curve (Ishii et al., 1990). Worms with mutations in *daf-2, mfn-1, cco-1, nuo-6*, or others genes had dramatically prolonged lifespan (Apfeld and Kenyon 1998; Ren et al., 2015), and their survival curves should be L (long-lived) patterned. Lifespan curves that are similar to the curve of N2 are N (normal) patterned. It is predictable that there exists a ST (steepened) patterned curve, for which the average lifespan is the same as that of N2, but its minimum and maximum lifespan is increased and decreased respectively. It seems that knocking down of *Y45F10D.4*, which encodes a putative iron-sulfur cluster assembly enzyme, is likely to produce the ST patterned curve (unpublished data). Unlike worms, whose body plan is simple, vertebrates have much more complicated tissues and organs and deletion of *clk-1* should be much more detrimental and unbearable for them, and this may explain the lethality phenotype observed in mutant mice.

By taking into account previously overlooked factors including individual specificity, the excessive response, and retardation of growth, here the reasons why *clk-1* RNAi worms have shortened while *clk-1* mutants have mildly prolonged or normal average lifespan was explained. We further categorized the lifespan curves into five patterns to indicate different kinds of lifespan variations reported in model organisms. The individual specificity in aging process, which is likely to be contributed by epigenetic factors, should be responsible for the variations of lifespan curves and deserves further investigations.

## Conflict of interest

The authors declare that there is no conflict of interest.

## Author contributions

Y.R. conceived this work; Y.R., Congjie Z., and W.G. performed the experiments; Y.R. and Chao Z. contributed to the analysis and interpretation of data; Y.R. and Chao Z. wrote the manuscript; Chao Z. provided reagents and instruments.

## Acknowledgements

*C. elegans* strains were provided by the Caenorhabditis Genetics Center (CGC), which is funded by NIH Office of Research Infrastructure Programs (P40 OD010440).

## Funding

This work was supported by the National Natural Science Foundation of China [Grant numbers 81200253 and 81570760]; the National Key Research and Development Program of China [Grant numbers 2016YFA0102200, 2017YFA0103900, and 2017YFA0103902]; One Thousand Youth Talents Program of China to C. Zhang; the Program for Professor of Special Appointment (Eastern Scholar) at Shanghai Institutions of Higher Learning [Grant number A11323]; the Shanghai Rising-Star Program [Grant number 15QA1403600]; and the Fundamental Research Funds for the Central Universities of Tongji University.

## References

Apfeld J, Kenyon C (1998) Cell nonautonomy of C. elegans daf-2 function in the regulation of diapause and life span. Cell 95(2):199–210

Brenner S (1974) The genetics of Caenorhabditis elegans. Genetics 77(1):71–94

Ewbank JJ, Barnes TM, Lakowski B, Lussier M, Bussey H, Hekimi S (1997) Structural and functional conservation of the Caenorhabditis elegans timing gene clk-1. Science 275(5302):980–3

Gu R, Zhang F, Chen G, Han C, Liu J, Ren Z, Zhu Y, Waddington JL, Zheng LT, Zhen X (2017) Clk1 deficiency promotes neuroinflammation and subsequent dopaminergic cell death through regulation of microglial metabolic reprogramming. Brain, behavior, and immunity 60:206–219

Hirsh D, Vanderslice R (1976) Temperature-sensitive developmental mutants of Caenorhabditis elegans. Developmental biology 49(1):220–35

Ishii N, Takahashi K, Tomita S, Keino T, Honda S, Yoshino K, Suzuki K (1990) A methyl viologen-sensitive mutant of the nematode Caenorhabditis elegans. Mutation research 237(3-4):165–71

Lakowski B, Hekimi S (1996) Determination of life-span in Caenorhabditis elegans by four clock genes. Science 272(5264):1010–3

Larsen PL, Clarke CF (2002) Extension of life-span in Caenorhabditis elegans by a diet lacking coenzyme Q. Science 295(5552):120–3

Nakai D, Yuasa S, Takahashi M, Shimizu T, Asaumi S, Isono K, Takao T, Suzuki Y, Kuroyanagi H, Hirokawa K et al. (2001) Mouse homologue of coq7/clk-1, longevity gene in Caenorhabditis elegans, is essential for coenzyme Q synthesis, maintenance of mitochondrial integrity, and neurogenesis. Biochemical and biophysical research communications 289(2):463–71

Ren Y, Chen S, Ma M, Yao X, Sun D, Li B, Lu J (2015) The activation of protein homeostasis protective mechanisms perhaps is not responsible for lifespan extension caused by deficiencies of mitochondrial proteins in C. elegans. Experimental gerontology 65:53–7

Ren Y, Yang S, Tan G, Ye W, Liu D, Qian X, Ding Z, Zhong Y, Zhang J, Jiang D et al. (2012) Reduction of mitoferrin results in abnormal development and extended lifespan in Caenorhabditis elegans. PloS one 7(1):e29666

Ren Y, Zhang C (2017) The excessive response: a preparation for harder conditions. Protein & cell

Shore DE, Carr CE, Ruvkun G (2012) Induction of cytoprotective pathways is central to the extension of lifespan conferred by multiple longevity pathways. PLoS genetics 8(7):e1002792

Simmer F, Moorman C, van der Linden AM, Kuijk E, van den Berghe PV, Kamath RS, Fraser AG, Ahringer J, Plasterk RH (2003) Genome-wide RNAi of C. elegans using the hypersensitive rrf-3 strain reveals novel gene functions. PLoS biology 1(1):E12

Takahashi M, Ogawara M, Shimizu T, Shirasawa T (2012) Restoration of the behavioral rates and lifespan in clk-1 mutant nematodes in response to exogenous coenzyme Q(10). Experimental gerontology 47(3):276–9

Vajo Z, King LM, Jonassen T, Wilkin DJ, Ho N, Munnich A, Clarke CF, Francomano CA (1999) Conservation of the Caenorhabditis elegans timing gene clk-1 from yeast to human: a gene required for ubiquinone biosynthesis with potential implications for aging. Mammalian genome: official journal of the International Mammalian Genome Society 10(10):1000–4

Zhang C, Wang K, Liu Z, Huang J, Zhou H, Lü J (2017) Effects of clk-1 on Lifespan in Caenorhabditis elegans. Chinese Journal of Cell Biology 39:745–755

